## Supplemental Figures S1-S5 for "Chromatin and gene expression changes during female Drosophila germline stem cell development illuminate the biology of highly potent stem cells"

### Supplemental Materials

#### Materials and Methods

Volume measurement

MTD-Gal4 > UASz-tdTomato flies with labeled germline cells are fed on wet yeast for 3 days. Ovaries are dissected carefully with forceps to break apart the surrounding muscle sheets. Dissected ovaries are fixed in 4% PFA for 10 min, stained with DAPI, and mounted in VECTASHIELD®. For *yl* mutant and control lines, dissected ovaries are stained with anti-vasa to mark germline cell volumes (see immunofluorescent staining method for details). Full z-stacked images of the ovary are taken under 40X oil objective with Leica TCS SP5 Confocal Microscope. Reconstruction of 3D images is performed with Oxford Instruments Imaris software. Volumes and total DAPI signal pixels of individual follicles are recorded and developmental stages of the follicles are manually noted. Growth rate changes are plotted and modeled with R.

Growth modeling

Local polynomial regression was used to accommodate for the complexity of the data. Smoothing factor =0.75 and degree of polynomial =2 were used. The data set is fairly dense with n>20 for each stage and is therefore suitable for local polynomial regression. However, the result of LOESS can be difficult to interpret because it cannot be written down as simple equations (Guthrie et al. 2002). To further test the developmental growth rate changes, multiple linear regression was applied with two conditional terms. Condition1 distinguishes follicles before (stage 1 to 5) and after (stage 6 to 10) nurse cell chromatin dispersion. Condition2 distinguishes follicles before (stage 1 to 7) and after (stage 8 to 10) onset of vitellogenesis. The model to be tested is as following:

log10 Volume = a0*log2DNA +a1*log2DNA*condition1 +a2*log2DNA*condition2

+b0+b1*condition 1 +b2*condition2

### Supplementary Tables

Table S1. All processed data

Table S2. E(z)-dependent gene downregulation.

Table S3. Mostly heterochromatic GSC genes with high H3K27ac

Table S4 Genes upregulated in young follicles relative to GSCs.

Table S5. 41 membrane transporters upregulated more than 10-fold in young follicles.

### Supplementary References

### Supplementary Figures

**Figure S1. Growth profiles (total germ cell volume) of Drosophila ovarian follicles**

**Figure S5. Analysis of chromatin changes at enhancers of Myc during oogenesis**

A. myc region chromatin accessibility assayed by ATACseq in GSCs and 4C-32C nurse cells (NC). B. Genomic DNA fragments from this region (1-11, indicated) cloned and integrated within Janelia GAL4 lines (Jennet et al. 2012) are shown. C. A summary of the GFP expression driven by 8 tested clones fragments. D. Enlarged view of GFP expression from clone 8 (R10G10), which is similar to clone 7 and is shown below. The pattern of expression is indicated to show low level (clones 7 and 8) and higher level (clones 9 and 10) GFP expression starting about stage 4.

Fig S1

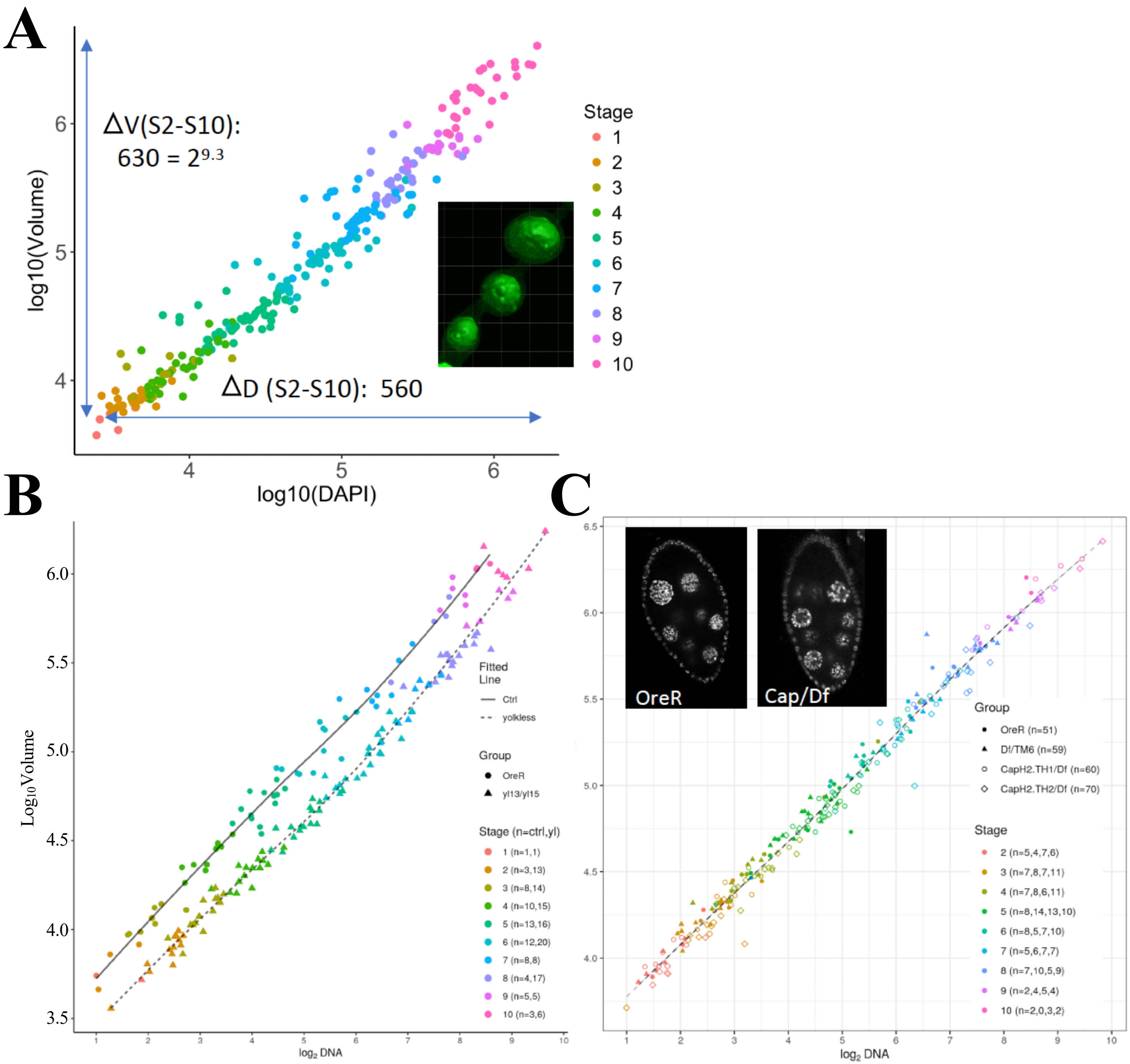
A
ΔV(S2-S10):
630 = 2^9.3
ΔD (S2-S10): 560
log10(Volume)
log10(DAPI)
Stage
1
2
3
4
5
6
7
8
9
10
B
Log10 Volume
log2 DNA
Fitted Line
Ctrl
yolkless
Group
OreR
yl13/yl15
Stage (n=ctrl,yl)
1 (n=1,1)
2 (n=3,13)
3 (n=8,14)
4 (n=10,15)
5 (n=13,16)
6 (n=12,20)
7 (n=8,8)
8 (n=4,17)
9 (n=5,5)
10 (n=3,6)
C
OreR
Cap/Df
log2 DNA
Group
OreR (n=51)
Df/TM6 (n=59)
CapH2.TH1/Df (n=60)
CapH2.TH2/Df (n=70)
Stage
2 (n=5,4,7,6)
3 (n=7,8,7,11)
4 (n=7,8,6,11)
5 (n=8,14,13,10)
6 (n=8,5,7,10)
7 (n=5,6,7,7)
8 (n=7,10,5,9)
9 (n=2,4,5,4)
10 (n=2,0,3,2)

**Fig S2**

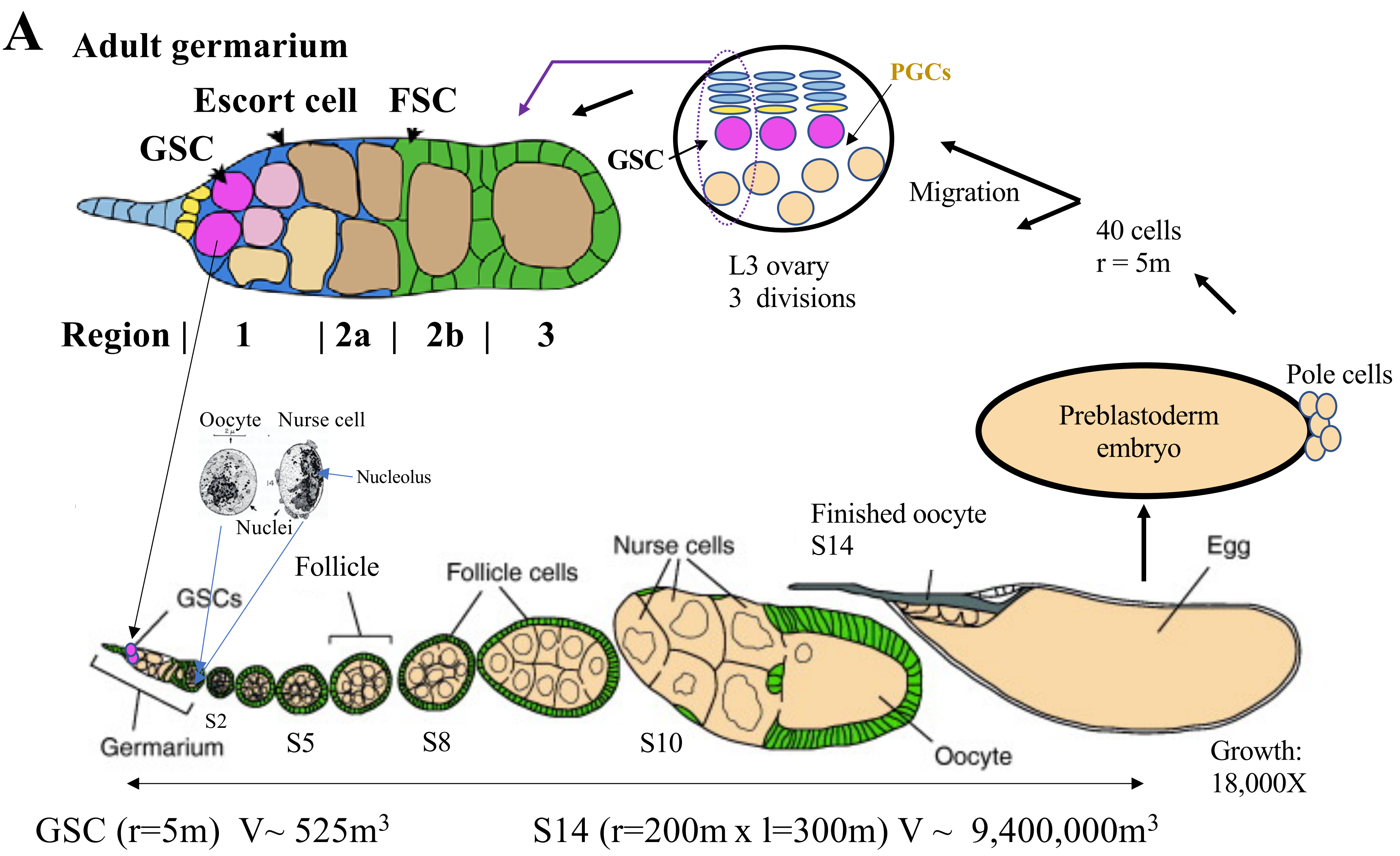
A
Adult germarium
Escort cell
FSC
GSC
Region | 1 | 2a | 2b | 3
PGCs
GSC
L3 ovary
3 divisions
Migration
40 cells
r = 5m
Pole cells
Preblastoderm embryo
Oocyte
Nurse cell
Nucleolus
Nuclei
Follicle
Follicle cells
Nurse cells
Finished oocyte
S14
Egg
GSCs
S2
Germarium
S5
S8
S10
Oocyte
Growth:
18,000X
GSC (r=5m) V~ 525m3
S14 (r=200m x l=300m) V ~ 9,400,000m3

**Fig S3**

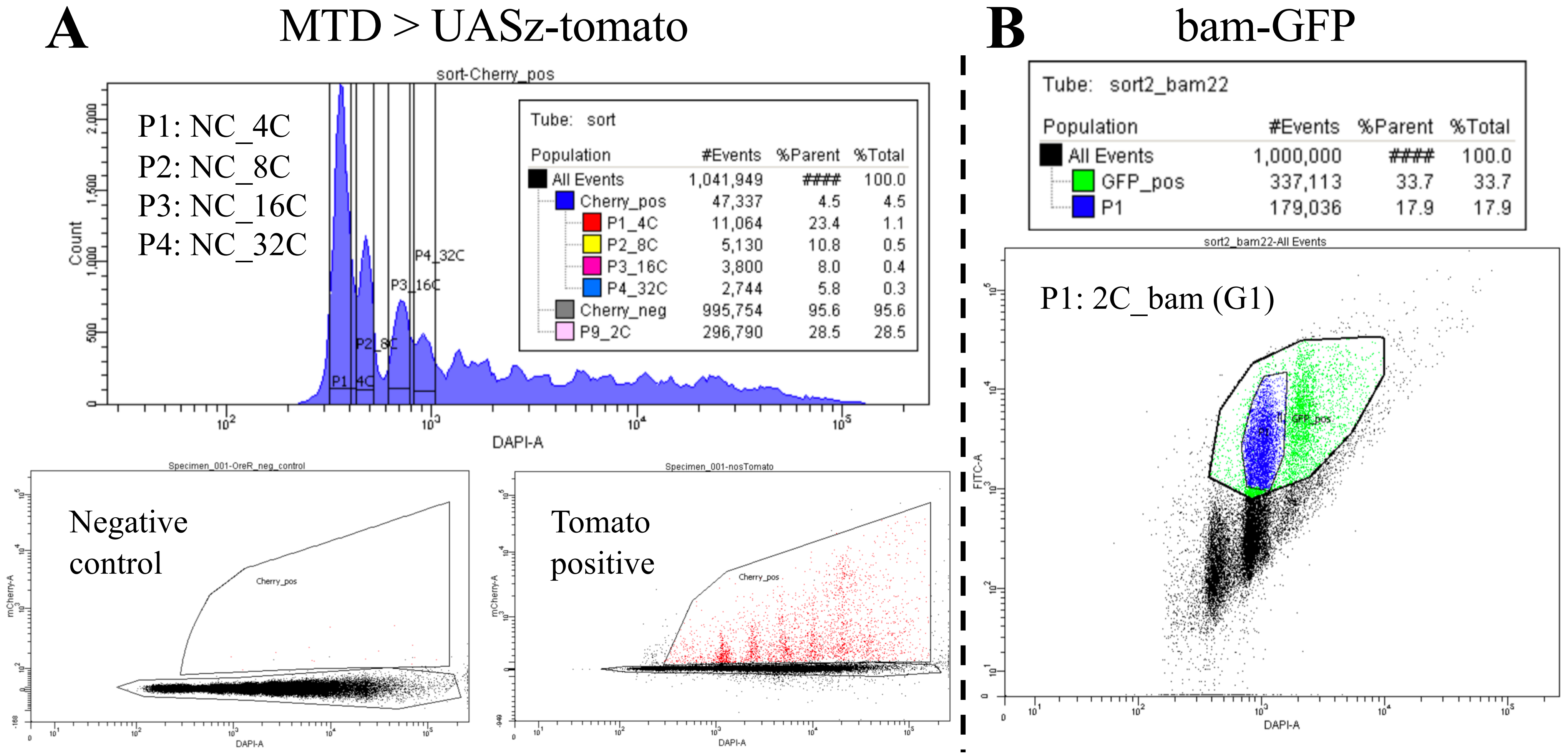
A
MTD > UASz-tomato
sort-Cherry_pos
P1: NC_4C
P2: NC_8C
P3: NC_16C
P4: NC_32C
Count
DAPI-A
Tube: sort
Population #Events %Parent %Total
All Events 1,041,949 #### 100.0
Cherry_pos 47,337 4.5 4.5
P1_4C 11,064 23.4 1.1
P2_8C 5,130 10.8 0.5
P3_16C 3,800 8.0 0.4
P4_32C 2,744 5.8 0.3
Cherry_neg 995,754 95.6 95.6
P9_2C 296,790 28.5 28.5
Specimen_001-OreR_neg_control
Negative control
Cherry_pos
mCherry-A
Specimen_001-nosTomato
Tomato positive
B
bam-GFP
Tube: sort2_bam22
Population #Events %Parent %Total
All Events 1,000,000 #### 100.0
GFP_pos 337,113 33.7 33.7
P1 179,036 17.9 17.9
sort2_bam22-All Events
P1: 2C_bam (G1)
FITC-A
DAPI-A

Fig S4

A

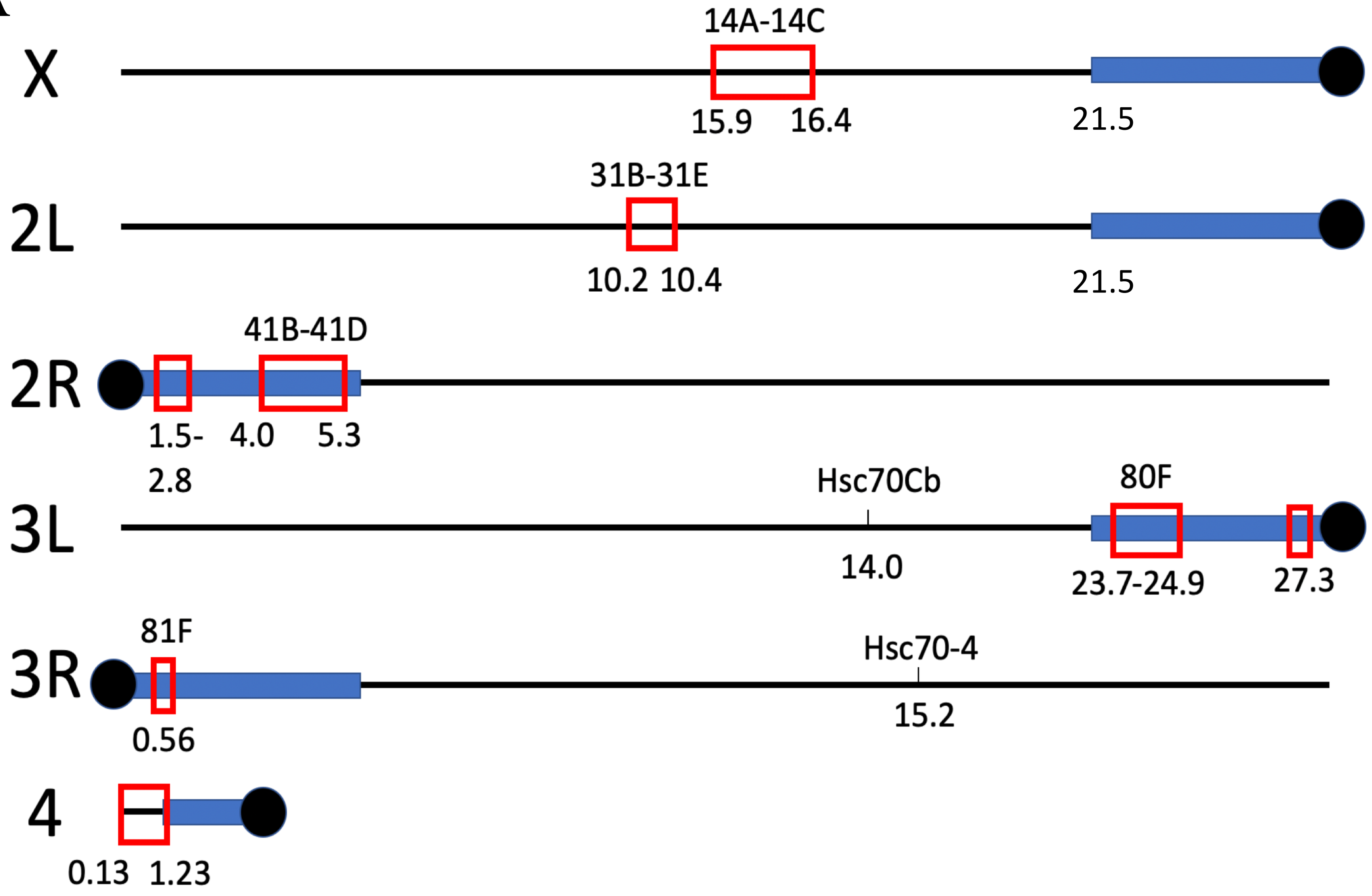
X
14A-14C
15.9
16.4
21.5
2L
31B-31E
10.2 10.4
21.5
2R
41B-41D
1.5-
2.8
4.0
5.3
3L
Hsc70Cb
14.0
80F
23.7-24.9
27.3
3R
81F
0.56
Hsc70-4
15.2
4
0.13
1.23

Fig S5

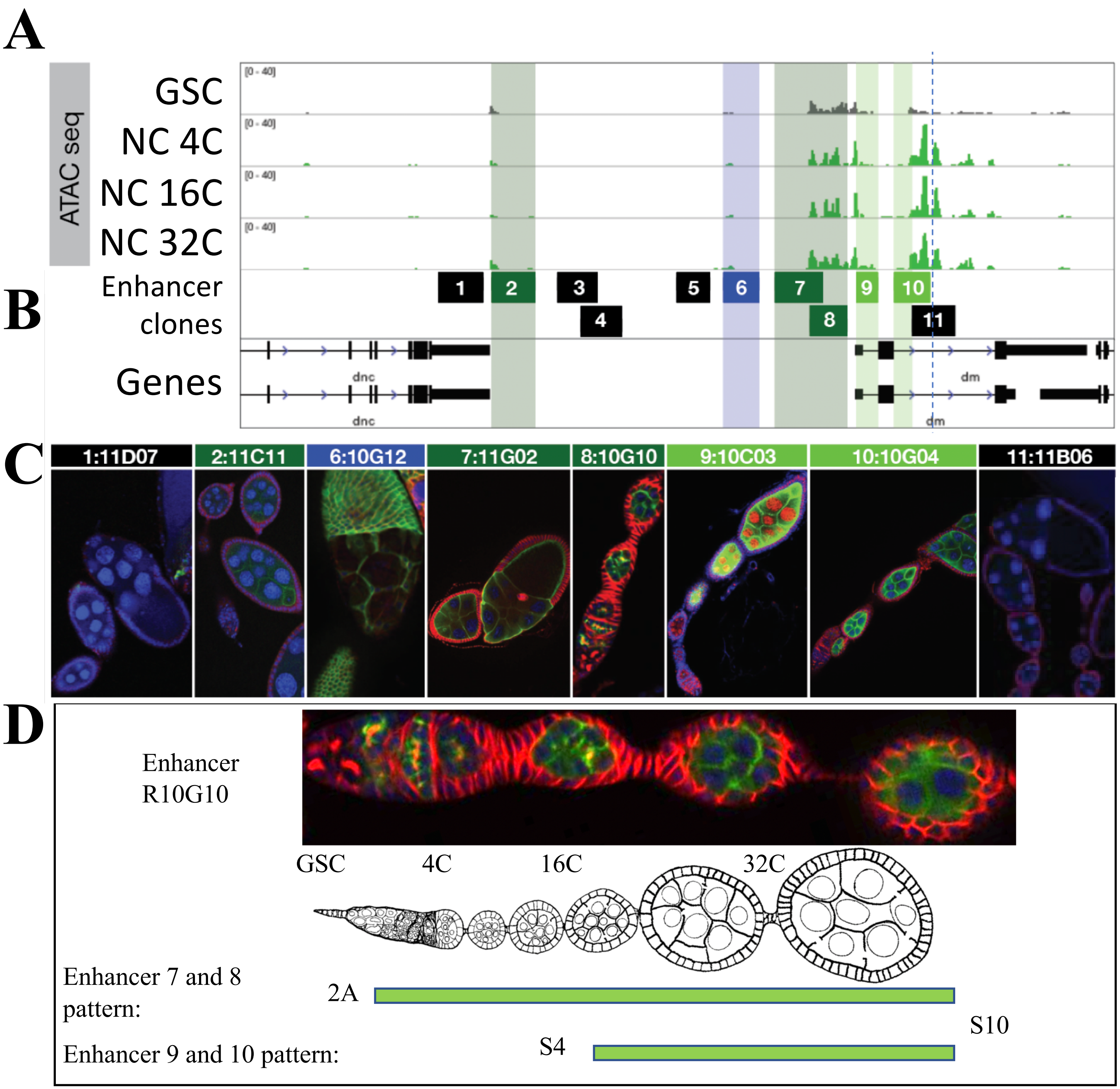
A
ATAC seq
GSC
NC 4C
NC 16C
NC 32C
B
Enhancer clones
1
2
3
4
5
6
7
8
9
10
11
Genes
dnc
dm
C
1:11D07
2:11C11
6:10G12
7:11G02
8:10G10
9:10C03
10:10G04
11:11B06
D
Enhancer R10G10
GSC
4C
16C
32C
Enhancer 7 and 8 pattern:
2A
S10
Enhancer 9 and 10 pattern:
S4
